# Mechanisms Coordinating Ribosomal Protein Gene Transcription in Response to Stress

**DOI:** 10.1101/2020.06.09.143263

**Authors:** Sevil Zencir, Daniel Dilg, Maria Paula Rueda, David Shore, Benjamin Albert

## Abstract

While expression of ribosomal protein genes (RPGs) in the budding yeast *Saccharomyces cerevisiae* has been extensively studied, a longstanding enigma persists regarding their co-regulation under fluctuating nutrient and stress conditions. Most (<90%) of RPG promoters display one of two distinct arrangements of a core set of transcription factors (TFs; Rap1, Fhl1 and Ifh1) and are further differentiated by the presence or absence of the HMGB box protein Hmo1. However, a third group of promoters appears not to bind any of these proteins, raising the question of how the whole suite of genes is co-regulated. We demonstrate that all RPGs are regulated by two distinct, but complementary mechanisms driven by the TFs Ifh1 and Sfp1, both of which are required for maximal expression in optimal conditions and coordinated down-regulation upon stress. At the majority of RPG promoters Ifh1-dependent regulation predominates, whereas Sfp1 plays the major role at all other genes. We also uncovered an unexpected, protein homeostasis-dependent binding property of Hmo1 at a large subset of RPG promoters. Finally, we show that the Ifh1 paralog Crf1, previously described as a transcriptional repressor, can act as a constitutive RPG activator in the W303 strain background when overexpressed. Our study thus provides a more complete picture of RPG regulation and may serve as a paradigm for unravelling RPG regulation in multicellular eukaryotes.

## Introduction

Ribosome biogenesis, one of the most energy consuming process in all organisms, is a major driver of rapid cell growth (Lempiainen and Shore, 2009; Warner, 1999). Ribosome production rates are estimated to be about 2-4,000 per minute in actively dividing *Saccharomyces cerevisiae* (hereafter yeast) and human cells, respectively. This intensive and complex production process involves several hundred Ribosome Biogenesis (RiBi) factors that direct the hierarchical assembly on to rRNA of the Ribosomal Proteins (RPs), which themselves constitute ∼50% of total protein copy number and ∼30% of protein mass in yeast. Regulated production of stoichiometric numbers of RPs is important in all organisms to maintain a functional proteome, as demonstrated by the discovery of specific mechanisms that prevent accumulation of unassembled RPs in both yeast and mammalian cells (Albert et al., 2019a; Lam et al., 2007; Sung et al., 2016a; Sung et al., 2016b; Tye et al., 2019). Impairment of ribosome biogenesis is also associated with cancer and ribosomopathies, a group of human diseases mainly caused by ribosomal protein haplo-insufficiency (Narla and Ebert, 2010). Understanding how cells produce roughly equimolar amounts of RPs in stress and non-stress conditions remains an open question of fundamental interest.

Most of our detailed knowledge on the expression of RPGs in eukaryotes is derived from studies of yeast, whose ribosomes contain 78 distinct RPs encoded by 138 RP genes (RPGs), of which 19 encode unique RPs while 118 encode duplicated RPs. RPGs are highly sensitive to gene dosage in *S. cerevisiae*, reflecting the need for RPG mRNA levels to be tightly controlled (Deutschbauer et al., 2005). Since most RPGs exist as duplicated copies, cells have developed specific regulations to achieve the required production of RPs in equimolar quantity (Parenteau et al., 2011). One of these mechanisms is encoded within their promoters by specific DNA sequences that facilitate or disfavour transcription, such that expression of single-copy RPGs is similar to that of the sum of duplicated paralogues (Zeevi et al., 2011). Pre-mRNA splicing, mRNA decay, translation and protein turnover provide additional layers of complexity that may act to equilibrate production of ribosome components (Petibon et al., 2016; Sung et al., 2016b; Warner, 1999).

Transcription of RPGs in yeast is tightly linked to cell growth, suggesting that TF binding at their promoters might be highly sensitive to stress and nutrient conditions. Consistent with this notion, inhibition of the Target Of Rapamycin Complex 1 (TORC1) kinase, a major transducer of nutrient signals, leads to cytoplasmic relocalization of growth-promoting transcription factor Sfp1 and release of the TF Ifh1 for RPG promoters (Albert et al., 2016; Cai et al., 2013; Downey et al., 2013; Jorgensen et al., 2004; Marion et al., 2004; Martin et al., 2004; Rudra et al., 2007; Rudra et al., 2005; Schawalder et al., 2004; Wade et al., 2004). Although, the stress-sensitive TF Sfp1 can increase or decrease RPG expression according to growth conditions, Sfp1 is mainly involved in regulation of RiBi genes (Albert et al., 2019b; Cipollina et al., 2008) whereas the transcription factor Ifh1 is defined as the main regulator for RPG transcription. Ifh1 binds almost exclusively to RPGs promoters (Schawalder et al., 2004), indicating that cells have developed a dedicated mechanism to regulate RPG expression. We showed previously that Ifh1 binding is directly linked to RNA Polymerase I (RNAPI) activity, and to levels of unassembled RPs in the nucleus, which allows cells to align RPG expression to both rRNA production and ribosome assembly, respectively (Albert et al., 2016; Albert et al., 2019a). The existence of these two mechanisms highlights the requirement for cells to develop specific processes for the maintenance of stoichiometric production of ribosome components.

How RPG co-regulation is achieved in *S. cerevisiae* still remains poorly understood since their promoters display a heterogeneous organization typically divided into two major categories (Category I and Category II, or Cat I and Cat II from hereon), both bound by the TFs Rap1, Fhl1 and Ifh1, and differentiated by the presence or absence of the High Mobility Group B (HMGB) protein Hmo1 (Hall et al., 2006; Jorgensen et al., 2004; Kasahara et al., 2007; Knight et al., 2014; Reja et al., 2015; Rudra et al., 2005; Schawalder et al., 2004; Wade et al., 2004). In addition, the existence of a third category of RPG promoters, apparently devoid of any transcription factors shared by the other two groups, constitutes one of the major obstacles in the complete understanding of mechanisms that allow RPG co-regulation. Only the general regulatory factor (GRF) Abf1, absent at other RPG promoters, is detected by chromatin immunoprecipitation (ChIP) at promoter of these genes (Bosio et al., 2017; Fermi et al., 2016). Consequently, it was suggested that these Cat III genes are expressed independently of the other RPGs (Reja et al., 2015), thus raising the question of how they could be co-regulated. Moreover, Crf1, a paralogue of Ifh1, has been reported to repress RPGs upon stress (Martin et al., 2004), but this mechanism seems not to operate in the widely used W303 strain background (Zhao et al., 2006).

In this study, we elucidate the common logic of the RPG regulatory network by evaluating both the architecture and activity of promoters under conditions of stress or modulation of TF levels. We uncovered an unexpected feature of the Ifh1 paralog, Crf1, which can act as a constitutively active version of Ifh1 in the commonly used W303 background, when overexpressed. Furthermore, we found that Hmo1 binding at RPGs is highly sensitive to proteotoxic stress. Importantly, we identified the TFs regulating the activity of the Category 3 promoters, which lack Rap1 binding. Our findings demonstrate that RPG co-regulation requires the complementary action of two different mechanisms, one involving Ifh1 and the other using Sfp1. The combination of these two mechanisms is required to rapidly coordinate the activity of the heterogeneously constituted RPG promoters upon stress.

## Results

### RPG promoter organization is heterogeneous but partitions into three distinct groups

Previous studies (Hall et al., 2006; Kasahara et al., 2007; Knight et al., 2014; Reja et al., 2015) have partitioned RPGs into several groups according to the arrangement of bound TFs upstream of their promoter, as described above and summarized in graphic form in **Figure 1A**. The GRF Rap1 and the TFs Ifh1, Fhl1, and Sfp1 all bind to Cat I and Cat II promoters, whereas only the HMGB protein Hmo1 binds uniquely at Cat I promoters (**Figure 1B**). Although Hmo1 binding constitutes one of the most striking distinguishing features of RPG promoters, whether or not Hmo1 is involved in regulation of RPG expression upon stress is not clear. Hmo1 binding at RPG promoters was reported to rapidly decrease after exposure to high temperature (Hall et al., 2006; Reja et al., 2015) but not following inhibition of the growth regulator TORC1, a finding that we confirm here (**Figure 1C**). Nevertheless, under both conditions (heat shock and rapamycin treatment) RPG expression is strongly repressed (**Figure 1D**). Furthermore, 20 minutes following inhibition of TORC1 by rapamycin treatment we find no evidence of Heat Shock Factor 1 (Hsf1) target gene activation, again linking Hmo1 to a thermal stress response (**Figure 1D)**. We recently showed that induced depletion of both topoisomerases 1 and 2 leads to the rapid induction of a proteotoxic stress pathway that we referred to as the Ribosome Assembly Stress Response (RASTR; ((Albert et al., 2019a)). During RASTR Hsf1 is activated and RPGs are down-regulated, as happens during heat shock (**Figure 1D**). Interestingly, Hmo1 binding levels rapidly decrease at Cat I RPG promoters following depletion of topoisomerases, despite the absence of thermal stress (**Figure 1C**). These data indicate that Hmo1 binding is affected by proteotoxic stress. However, the fact that Hmo1 binding is not totally abolished upon proteotoxic stress, and that Hmo1 is not released from RPG promoters during TORC1 inactivation, clearly demonstrates that release of Hmo1 from promoters during stress is not a prerequisite for RPGs downregulation, at least upon TORC1 inactivation. In contrast, Ifh1, an essential activator dedicated almost exclusively to RPG transcription, is rapidly released from promoters upon a wide range of stresses, including nutrient starvation, heat shock or TORC1 inactivation (Albert et al., 2016; Cai et al., 2013; Downey et al., 2013; Jorgensen et al., 2004; Martin et al., 2004; Rudra et al., 2007; Rudra et al., 2005; Schawalder et al., 2004; Wade et al., 2004). All promoters from Cat I and II have Rap1, Fhl1, Sfp1, Ifh1 at their promoters, but the thirteen Cat III promoters display a very different organization. This group of promoters is bound by Abf1 and depleted for Rap1, Fhl1, Ifh1, and Sfp1, apart from both *RPL1A* and *RPL18B*, whose promoters are bound by Rap1 but fail to recruit Fhl1, Ifh1 or Sfp1 at detectable levels (Kasahara et al., 2007; Knight et al., 2014; Reja et al., 2015; Schawalder et al., 2004)). Cat III genes are expressed at similar levels to other RPGs (**Sup Figure 1**) and code for both large and small subunit (60S and 40S) ribosomal proteins (**Figure 1A**). However, the absence of common transcription factors shared with other RPGs raises the question of how they are co-regulated with Cat I and Cat II genes. A previous study reported the presence of an Fhl1 binding motif in close proximity to Abf1 binding sites at several Cat III genes (Fermi et al., 2016). Nevertheless, the *in vivo* functionality of these Fhl1 binding sites remains controversial since Fhl1’s ability to bind these sequences has only been confirmed by *in vitro* experiments and ChIP experiments, by us and others, have failed to detect significant Fhl1/Ifh1 binding at Cat III promoters (**Figure 1B**; (Fermi et al., 2016; Reja et al., 2015,Knight, 2014 #23,Cai, 2013 #61)).

**Figure 1.**
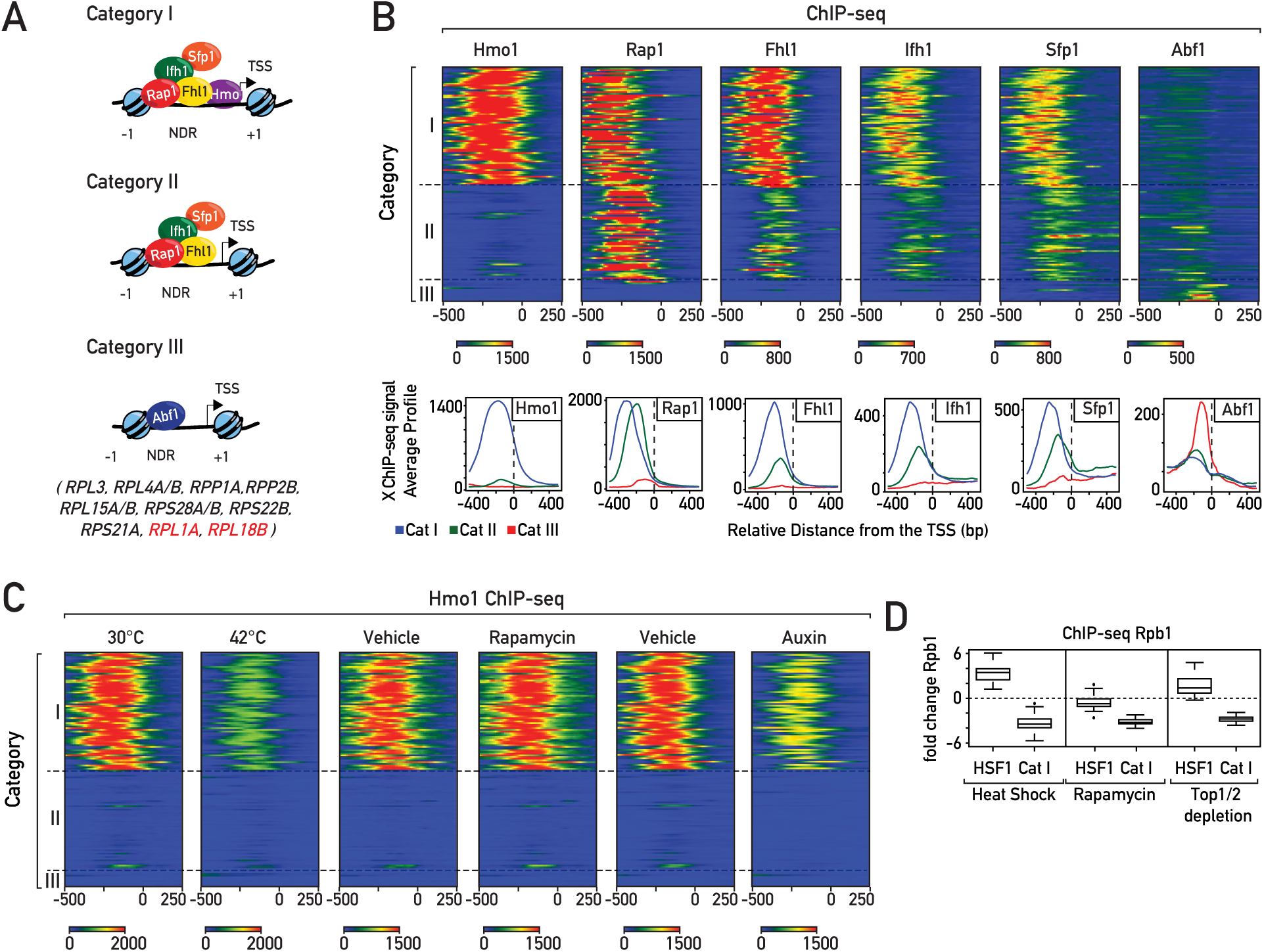
Heterogeneous organization of RPG promoters. **(A)** Schematic representation of RPG categories according to their promoter nucleosome and transcription factor architecture. In each schema the left-most nucleosome represents the first stable −1 nucleosome, the right-most nucleosome represents the +1 nucleosome, black arrows represent the Transcription Start Site (TSS), and NDR corresponds to the nucleosome-depleted region (Hall et al., 2006; Kasahara et al., 2007; Knight et al., 2014; Reja et al., 2015; Schawalder et al., 2004; Wade et al., 2004). Genes included in Cat III (Knight et al., 2014) are reported. *RPL1A* and *RPL18B* are two peculiar cases in this group since Rap1 is detected at their promoters, though not Ifh1 and Fhl1. **(B)** Heat maps showing ChIP-seq signals for transcription factors Hmo1, Rap1, Ifh1, Sfp1 and Abf1 (from left panel to right panel) at RPG Categories I, II and III. Signals for a window of −500 to +250 bp relative to the dyad axis of the +1 nucleosome (0) are displayed (X-axis). **(C)** Heat maps showing Hmo1 ChIP-seq signals at RPG promoters in cells following either 5 min of heat shock at 42°C, 20 min treatment with or without rapamycin, or rapid depletion of Top1/2 by 20 min of auxin treatment. Signals for a window of −500 to +250 bp relative to the dyad axis of the +1 nucleosome (0) are displayed (x-axis). **(D)** Box plots of log2 RNAPII (Rpb1) ChIP-seq change at Hsf1 target genes and Category I RPG promoters following 5 min of heat shock at 42°C (left panel), 20 min of rapamycin treatment (middle panel), or Top1/2 depletion (right panel).

### Coregulation of the three groups of RPGs is adjusted according to growth conditions

To gain insight into the extent to which expression of the three categories of RPGs is coordinated, we first measured their transcription following TORC1 inactivation by rapamycin, which mimics nutrient starvation (e.g. carbon, nitrogen, phosphate or amino acid limitation). Importantly, inactivation of TORC1 also occurs in other types of stress, such as osmotic stress, redox stress, or caffeine treatment. We used RNAPII ChIP-seq as proxy for transcription since steady-state mRNA levels can mask transcription effects that are buffered by compensatory mRNA stability changes (Sun et al., 2013). As expected, inhibition of growth by rapamycin triggers global changes in the transcription program (**Figure 2A**), and more specifically leads to rapid downregulation of the three groups of RPGs, at both 5 and 20 min post treatment, without significant differences between Cat I, II, and III genes (**Figure 2A**, left and middle panels). Moreover, RiBi genes, known to be regulated by Sfp1 according to growth conditions (Cipollina et al., 2005; Cipollina et al., 2008), are also downregulated to a similar extent, though in a more heterogeneous fashion (**Figure 2A**). These findings reveal that, despite heterogeneous promoter structures, cells have developed mechanisms of comparable efficiency to rapidly coregulate RNAPII recruitment to all RPGs upon TORC1 inactivation. Nevertheless, we noted that following a 1 hr rapamycin treatment Cat III genes (largely bound by Abf1) are less downregulated and display a pattern more similar to RiBi genes than to the other RPGs (**Figure 2A**, right panel). This latter result points to the existence of distinct mechanisms to regulate the three groups of RPGs, allowing the cell to differentially modulate expression of RPGs under certain conditions.

**Figure 2.**
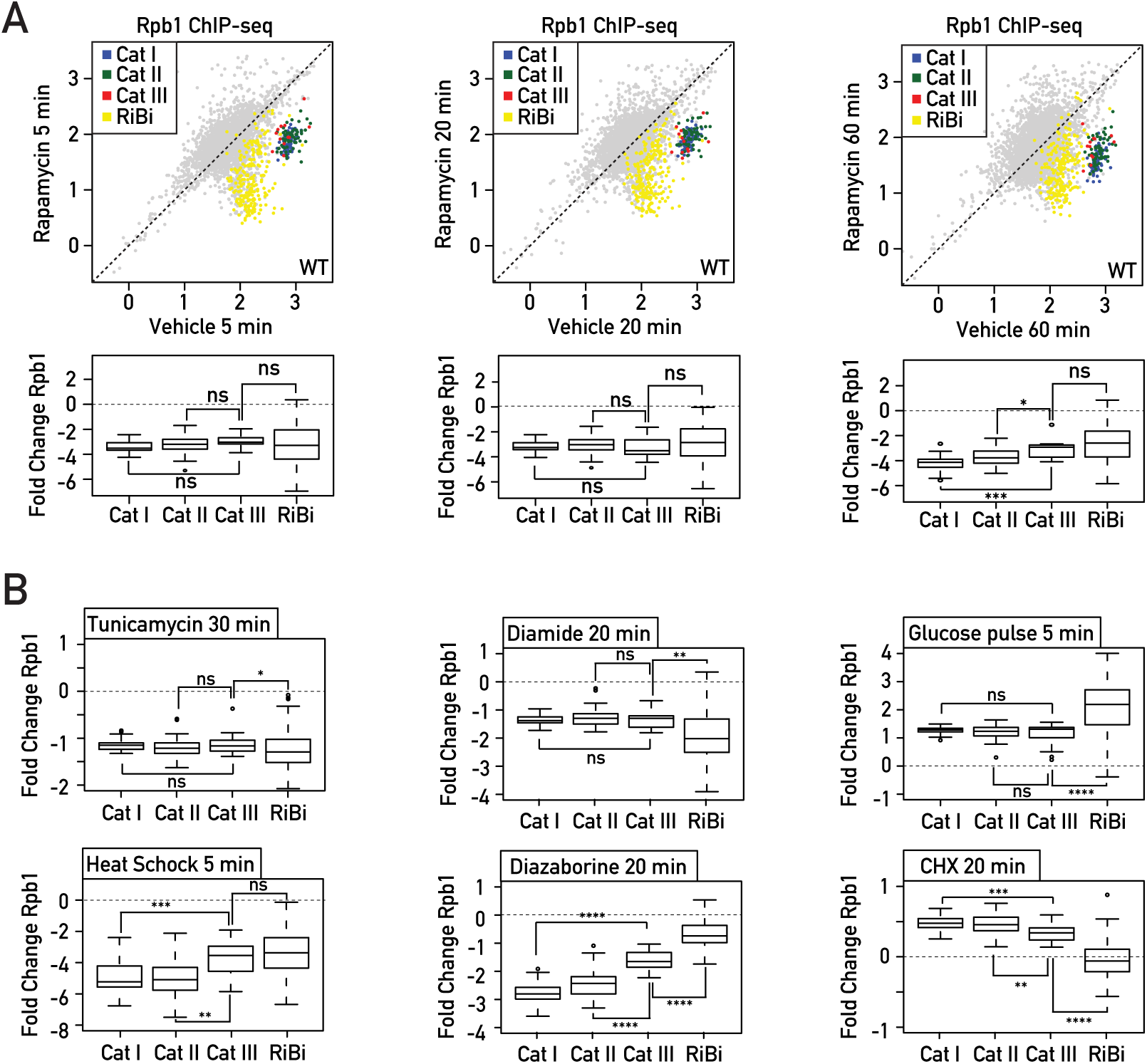
The three distinct categories of RPGs are co-regulated according to growth conditions. **(A)** Scatter plots (top panels) comparing RNAPII binding (as measured by Rpb1 ChIP-seq) in WT cells treated with rapamycin (Y-axis) or vehicle (X-axis) for 5 min (left panel), 20 min (middle panel) and 60 min (right panel). Each dot represents a gene (5041 in total) and genes are color-coded according to functional groups as Cat I (blue), Cat II (green) and Cat III (red) RPGs; RiBi genes (yellow); all other genes (grey). For RNAPII, the average signal was quantified from the TSS to the transcription termination site (TTS). The scale for both the X-axis and the Y-axis is log10. Bottom panels display the corresponding box plots for the four indicated gene categories. **(B)** Box plots showing RNAPII (Rpb1) ChIP-seq change for RPGs and RiBi genes in different growth conditions; upper panel shows the result of treatment with tunicamycin (30 min, left panel), diamide (20 min, middle panel) and glucose pulse (5 min, right panel); bottom panel shows the result of treatment with heat shock (5 min, left panel), diazaborine (20 min, middle panel) and cycloheximide (CHX, 20 min, right panel). Asterisks show significant difference according to the Mann-Whitney test (*: p<0.05, **: p<0.01, ***: p<0.001, ns: not significant).

To pursue this observation further we examined other conditions in which RPG transcription is known to be strongly affected, including glucose addition to cells growing in a poor carbon source (glycerol), heat shock, and oxidative stress (diamide treatment). We also tested the effects of arresting the secretion pathway, blocking translation elongation and inhibiting ribosome biogenesis by treating cells with tunicamycin, cycloheximide, and diazaborine, respectively. Interestingly, downregulation following tunicamycin or diamide treatments and upregulation after a 5 min glucose pulse are very similar for the three RPG categories, whereas heat shock, arrest of ribosome biogenesis by diazaborine or cycloheximide treatment all trigger a more heterogeneous response, with Cat III genes responding more weakly to heat shock and diazaborine compared to Cat I and II genes, but more strongly to cycloheximide (**Figure 2B**). These results demonstrate that strict RPG co-regulation is operative in some but not all conditions, suggesting that the different categories of RPGs are regulated through at least partly distinct processes.

### Co-regulation of RPGs through Ifh1-dependent and Ifh1-independent mechanisms

It was previously reported that the expression of RPGs is aligned to RNAPI activity (Albert et al., 2016; Laferte et al., 2006). To test whether all three categories of RPGs have this ability for co-regulation with RNAPI, we measured their expression following TORC1 inactivation by rapamycin treatment in cells where RNAPI is rendered constitutively active by expression of its Rrn3 and Rpa43 subunits as a fusion protein (CARA strain: Constitutive Association of Rrn3 and Rpa43; (Laferte et al., 2006)). As expected, transcriptional regulation of RPGs upon stress was specifically affected in the CARA strain, where RNAPI inhibition is blocked, whereas expression of other TORC1-sensitive genes remained unchanged (**Figure 3A**, compare with **Figure 2A**, middle panels). Contrary to what we observed previously in a wild-type strain, the transcriptional response of the three RPG categories was not homogeneous after 20 minutes of rapamycin treatment in the CARA strain. Cat I and II genes, which have Ifh1 at their promoters, were very weakly downregulated, whereas Cat III genes, whose promoters are devoid of Ifh1, were repressed more strongly (**Figure 3A-B**). This observation is consistent with our previous finding that the crosstalk between RNAPI and RPG expression, which is abrogated in the CARA strain, is dependent upon Ifh1 and its interaction with the CK2-Utp22-Rrp7-Ifh1 (CURI) complex, which is itself responsive to RNAPI activity (Albert et al., 2016). However, we noted that repression of Cat III genes upon TORC1 inactivation in the CARA strain was still not complete (**Figure 3B**), suggesting that Ifh1 might play a partial role in the regulation of these genes in coordination with RNAPI activity.

**Figure 3.**
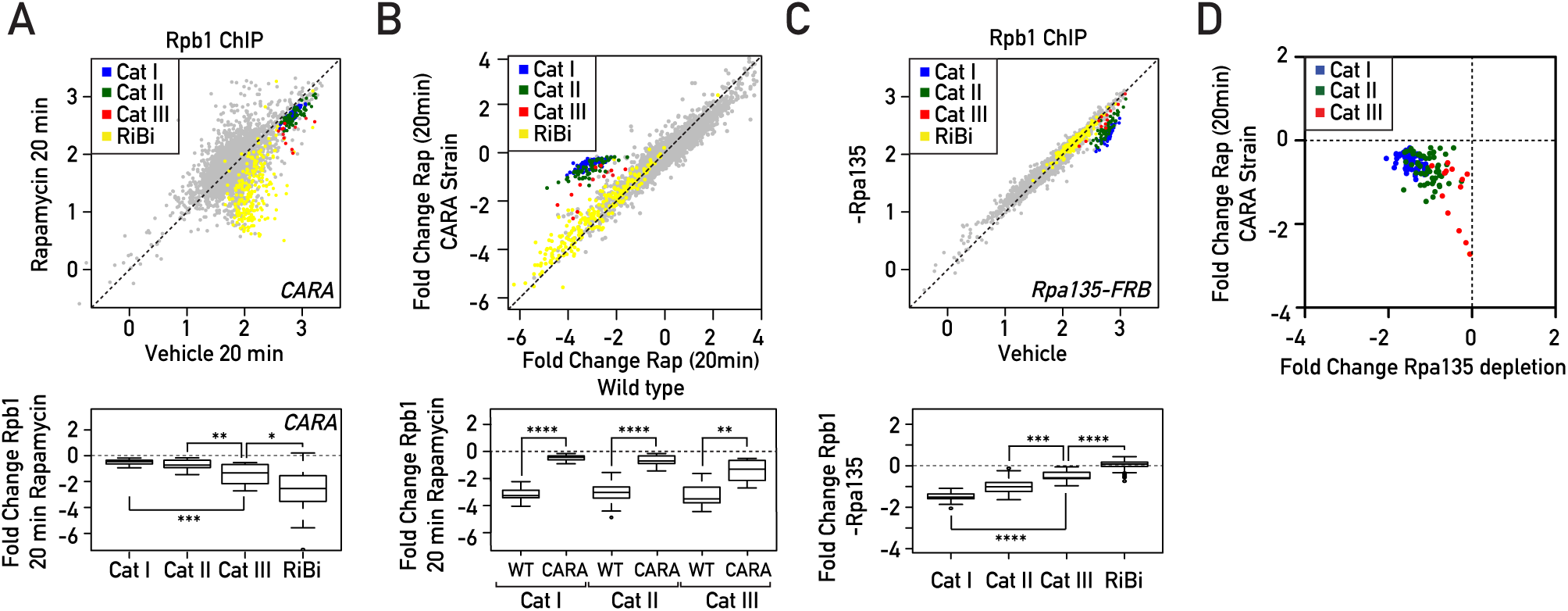
Coordinated downregulation of RPGs expression is carried out by Ifh1-dependent or Ifh1-independent processes. **(A)** RNAPII ChIP-seq in CARA strain cells (Y-axis) versus WT cells (X-axis) following to treatment with Rapamycin for 20 min. Bottom panels display the corresponding box plots for the four indicated gene categories. **(B)** Fold change of RNAPII binding following 20 minutes of rapamycin treatment in CARA strain cells (Y-axis) versus fold change of RNAPII binding following 20 minutes of rapamycin treatment in WT cells (X-axis). Bottom panels display the corresponding box plots for the three indicated gene categories. **(C)** RNAPII ChIP-seq in Rpa135 nuclear-depleted cells (−Rpa135; Y-axis) versus non-depleted cells (Vehicle; X-axis). Bottom panels display the corresponding box plots for the four indicated gene categories. **(D)** Scatter plots comparing RNAPII (Rpb1) binding fold change at RPGs categories in CARA strain cells treated with Rapamycin for 20 min (Y-axis) versus Rpa135 nuclear-depleted cells. Genes are color-coded according to functional groups: Cat I (blue), Cat II (green) and Cat III (red) RPGs.

To further test the ability of Cat III gene expression to be influenced by RNAPI activity, we chose to decrease rRNA production by nuclear depletion of one subunit of RNAPI (Rpa135), using the anchor away-system (Haruki et al., 2008). Remarkably, 60 minutes of Rpa135 nuclear depletion led to specific downregulation of RPG expression whilst other groups of genes, such as RiBi genes (used here as a proxy of TORC1 activity), were unaffected (**Figure 3C**). We confirmed that all RPGs, including Cat III genes, are sensitive to a decrease of RNAPI activity, but to different extents. These results also confirm that several mechanisms co-regulate RPGs following TORC1 inactivation; one is Ifh1/RNAPI-dependent and only partially effective on Cat III genes, while acting strongly on Cat I and II genes. Another mechanism is Ifh1/RNAPI-independent and strongly affects Cat III genes, but only weakly influences Cat I and II genes. The complementary action of these two mechanisms appears to be required to fine-tune co-regulation of all RPGs following TORC1 inactivation. Consistent with this notion, RPGs that were heavily affected by an Ifh1/RNAPI-independent process (Cat III genes) were the least affected by the Ifh1-dependent process (**Figure 3D**).

### Crf1 is a non-regulatable activator of RPGs

It was initially suggested that Crf1, an Ifh1 paralog, acts as a negative regulator of RPGs following TORC1 inactivation by competing with Ifh1 for binding to RPG promoters (Martin et al., 2004). Indeed, Crf1 has a well conserved forkhead-associated binding (FHB) domain (**Figure 4A**) through which it binds to Fhl1 (Martin et al., 2004), and deletion of *CRF1* in the TB50 strain background reduces RPG downregulation in response to rapamycin treatment (Martin et al., 2004; Zhao et al., 2006). Surprisingly, though, deletion of *CRF1* does not prevent the downregulation of RPGs following rapamycin treatment in the W303 genetic background (Zhao et al., 2006), where Crf1 may be weak or non-existent, as judged by western blot or RNAPII binding at its ORF (data not shown). In order to reveal further structural properties of different RPG categories, we tested the idea that forced Crf1 expression might inhibit growth in the W303 background through competition with Ifh1 for binding at RPG promoters by introducing a plasmid bearing *CRF1* coding region under control of the strong and constitutive *PGK1*. Surprisingly, overexpression of Crf1 had no effect on growth under optimal conditions, but instead suppressed the growth defect of different hypomorphic alleles of *IFH1* (*ifh1-AA, Ifh1-s, ifh1-6*; **Figure 4B**) and, remarkably, rescued the lethality of *IFH1* deletion (**Figure 4C**).

**Figure 4.**
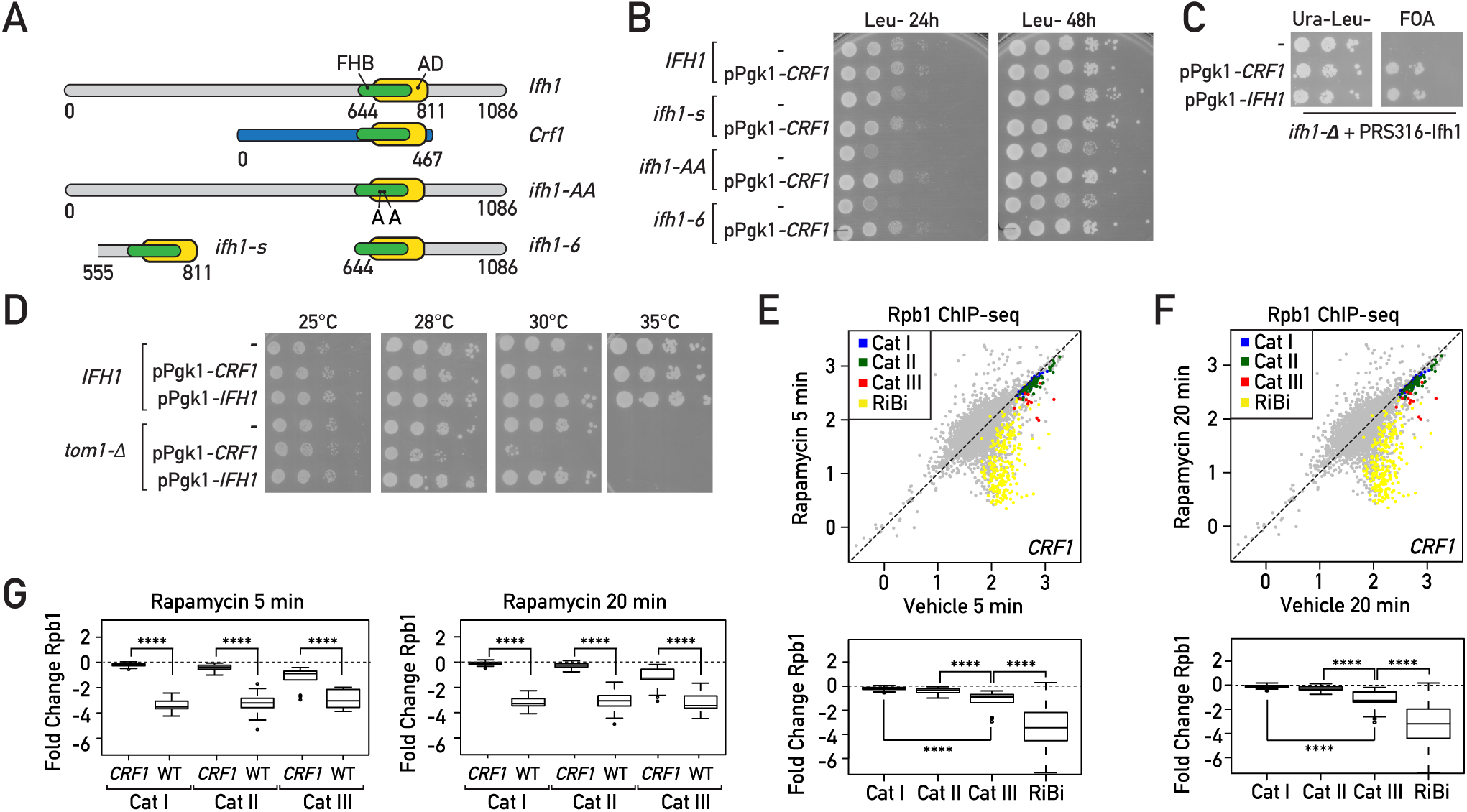
Crf1 is a non-regulatable activator of RPGs. **(A)** Schematic representation of conserved domains in full length Ifh1, Crf1 proteins and a series of Ifh1 points mutated S680A/S681A (*ifh1-AA*) or truncated alleles; Ifh1 mutant that removed all sequences upstream of the FHB and linked activation domain (*ifh1-6*); an extremely short version of Ifh1 (*ifh1-s*) containing essentially the FHB and downstream activation domain (from top panel to bottom panel). FHB: Fork Head Binding domain, AD: Activation domain. **(B)** 10-fold serial dilution of wild-type (*IFH1*) or hypomorphic alleles of Ifh1 *(ifh1-s, ifh1-AA, Ifh1-6*), transformed with a plasmid bearing CRF1 coding region under the control of PGK promoter (pPGK1-CRF1) or with the empty pPGK1 vector (−). Cells were spotted onto the indicated selective media and the plates were incubated at 30°C for 24hr or 48 hr. **(C)** A null allele (*ifh1-Δ*) of haploid tester strain is complemented by the corresponding WT genes borne on *URA3*-containing plasmids (pRS316-Ifh1). Complementation is tested by examining whether plasmids expressing Crf1 (pPGK1-CRF1) or Ifh1 (pPGK1-CRF1) from a strong promoter bypass the lethal phenotype, and monitored by growth on FOA. The empty pPGK1 vector (−) is used as negative control. Plates without FOA (Ura−Leu−) are used as control to confirm that the same number of cells were spotted. Plates were incubated at 30°C for 48 hr. **(D)** 10-fold serial dilution of wild-type (*IFH1*) or *TOM1* deleted cells *(tom1-Δ*), transformed with a plasmid bearing *CRF1* or *IFH1* coding regions under the control of PGK promoter (pPGK1-CRF1 or pPGK1-IFH1, respectively) or with the empty pPGK1 vector (−). Cells were spotted onto the indicated selective media and the plates were incubated at different temperatures for 48hr. (**E, F**) Scatter plots comparing RNAPII (Rpb1) ChIP-seq in Crf1 expressing cells after 5 min (Y-axis, **E**) or 20 min (Y-axis, **F**) Rapamycin treatment to non-treated cells (Vehicle; X-axis). Bottom panels display the corresponding box plots for the four indicated gene categories. Gene groups are color-coded as indicated above. (**G**) Box plots comparing RNAPII (Rpb1) binding fold change at RPGs categories in *CRF1* expressing and WT cells treated with rapamycin for 5 min (left panel) or 20 min (right panel). Asterisks show significant difference according to Mann-Whitney test.

Although Crf1 contains an FHB domain highly similar to that of Ifh1, it completely lacks the C- and N-terminal regions of Ifh1 that are implicated in the removal of Ifh1 from RPG promoters upon stress (**Figure 4A**; (Albert et al., 2016; Albert et al., 2019a)). The absence of these C- and N-terminal extensions may help Crf1 to compete efficiently with Ifh1 upon stress. Interestingly, elevated *CRF1* protein level becomes highly toxic at 30°C in a strain deleted for *TOM1*, which encodes a ubiquitin ligase required for degradation of ribosomal protein produced in excess, suggesting that Crf1 dysregulates RP production (**Figure 4D**). Consistent with this idea, increased *CRF1* expression prevents the repression of Cat I and II genes at both 5 and 20 minutes following rapamycin treatment (**Figure 4E-4F**; compare with **Figure 2A**, left and middle panels). For the case of Cat III, increased *CRF1* expression has a smaller but still highly significant effect (**Figure 4E-4F**) in comparison to other genes downregulated following rapamycin treatment, such as RiBi genes, which are completely unaffected by *CRF1* expression. This latter finding bolsters the idea that Fhl1 binding sites in proximity to Abf1 (Fermi et al., 2016) can act, through the FHA-FHB interaction, to recruit either Crf1 or Ifh1. Nevertheless, as shown above, a major part of the transcriptional downregulation of Cat III genes is carried out by an Ifh1-independent process.

### Both Sfp1 and Ifh1 are recruited to Cat III promoters

We next examined DNA sequence features of Cat III promoters in order to identify specific regulators of this group of genes. This analysis revealed that RPG promoter categories are distinguished not just by transcription factor binding heterogeneity, but also by nucleosome depleted region (NDR) size and G/C content (**Figure 5A**). These features are distinctive between the three categories of RPGs: Cat I promoters have a high G/C content and the largest NDRs, whilst Cat III promoters have the smallest NDRs and the lowest G/C content (**Figure 5B**). The G/C content of Cat I promoters may be important to maintain a large NDR, by promoting RSC (Remodeling the Structure of Chromatin) and Hmo1 binding (Badis et al., 2008; Knight et al., 2014; Kubik et al., 2018). On the other hand, it could also introduce a bias in ChIP assays, due to the strong bias in formaldehyde crosslinking for G/C base pairs (Rossi et al., 2018), which might lead to low detection of proteins such as Abf1, which is enriched at Cat III promoters (Fermi et al., 2016). We also noted that Cat III promoters are enriched in a motif (gAAAATTTTc) bound by Sfp1, both *in vitro* and *in vivo* (**Figure 5C**; (Albert et al., 2019b; Zhu et al., 2009)), which is itself largely depleted for G/C base pairs. We recently reported (Albert et al., 2019b) that Sfp1 binding is undetectable by ChIP at RiBi promoters enriched for the Sfp1 binding motif but can be revealed only by an alternative assay that does not require formaldehyde cross linking, Chromatin Endogenous Cleavage (ChEC; (Schmid et al., 2004)). We thus used published ChEC-seq data (Albert et al., 2019b) to examine binding of Sfp1 at RPGs promoters. Strikingly, Sfp1 ChEC-seq signal reveals a high level of binding at Cat III promoters, comparable to the signals observed at RiBi genes, and more globally shows an opposite binding pattern at promoters of the three RPG categories compared when comparing it to the Sfp1 signal generated by ChIP (**Figure 5D, 5E**). This result strongly suggests that Sfp1 could be the missing regulator of Abf1-dependent genes following TORC1 inactivation, and that structural properties of Cat III promoters probably limit its ability to be ChIPed at these sites. Interestingly, Ifh1 ChEC-seq also revealed significant binding at Cat III promoters, where it is essentially undetectable by ChIP, in comparison with RiBi gene promoters or a group of two hundred RNAPII promoters chosen at random (**Figure 5D, 5E**). However, contrary to Sfp1 ChEC-seq, Ifh1 ChEC-seq yielded a stronger signal on Cat I and Cat II compared to Cat III promoters. This Ifh1 binding pattern revealed by ChEC-seq is fully consistent with results presented above suggesting that Ifh1 has a minor but significant role in the regulation of Cat III genes in comparison to its major role at Cat I and II genes.

**Figure 5.**
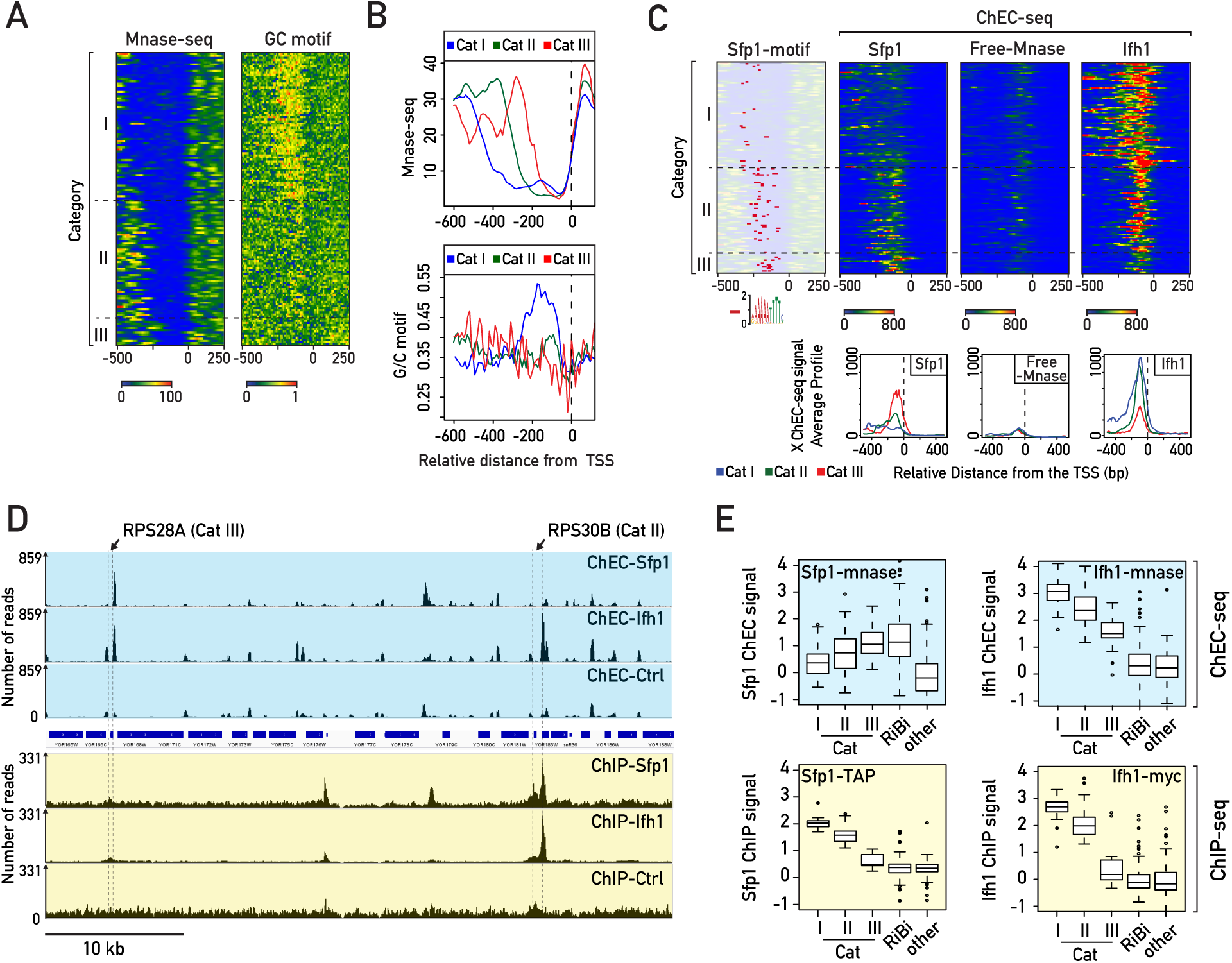
Sfp1 and Ifh1 are directly recruited at Cat III promoters. **(A)** Heat maps showing MNase digestion patterns (left panel) and G/C content (right panel) at RPGs promoters. Signals for a window of −500 to +250 bp relative to the +1 nucleosome (0) are displayed (X-axis). **(B)** Average plots of MNase digestion patterns (upper panel) and G/C content (lower panel) at three categories of RPGs. **(C)** Heat maps showing Sfp1-binding motif and Sfp1 ChEC-seq signal for 150 sec of Ca^+2^ treatment, or Ifh1 ChEC-seq signal after 150 sec of Ca^+2^ treatment at the indicated RPGs promoters. Control for ChEC-seq signal (free-MNase, 20 min following Ca^+2^ addition) is used as background control. Average plots of Ifh1, Sfp1, Free-Mnase ChEC-seq signal at three categories of RPGs is also shown (Lower panel). **(D)** Genome browser tracks comparing Sfp1-MNase, Ifh1-MNase and free MNase ChEC-seq signals (blue background) to Sfp1-TAP, Ifh1-Myc and untagged ChIP-seq read counts (yellow background) at *RPS28B* (Cat I) and *RPS30B* (Cat II) RPGs. The position of indicated RPGs ORF are shown above of the tracks. **(E)** Box plots of log2 ChEC-seq or ChIP-seq signal related to ChEC or ChIP control (free MNase or untagged strains) at promoters of different groups of genes [Cat I, II, III, ribosome biogenesis (RiBi) genes, others (200 randomly chosen protein-coding genes)] for Sfp1-MNase and Ifh1-MNase (ChEC-seq, blue background) or Sfp1-TAP and Ifh1-Myc (ChIP-seq, yellow background), respectively.

### Coordinated downregulation of RPGs by complementary regulation of both Sfp1 and Ifh1

Taken together, the results described above allow us to propose a new organizational principle of TFs at RPG promoters in which Ifh1 and Sfp1 can bind to and influence expression of all RPGs, including the small group of Cat III genes bound by the GRF Abf1, instead of Rap1. In order to challenge this model by a functional assay, we measured RNAPII recruitment in the absence of factors detected at the Cat III promoters: Abf1, Sfp1, Ifh1. Abf1 depletion triggered a very slight decrease of RNAPII recruitment at Cat III genes with only *RPL4A* being strongly affected (**Figure 6A**). In contrast, numerous non-RP Abf1 target genes were strongly affected by Abf1 depletion across the genome (**Supplemental Figure S6A**). Interestingly, these other Abf1 target genes are unaffected by TORC1 inactivation, suggesting that the transcriptional effect observed following TORC1 inactivation at Category III RPGs is independent of a decrease in Abf1 binding at these promoters. Next, we assessed the consequence of depletion of stress sensitive factors Ifh1 and Sfp1. Consistent with our previous results (Albert et al., 2019b), Ifh1 depletion triggered strong downregulation of Rap1-dependent genes and a significant though smaller decrease in transcription of Cat III genes (**Figure 6B**). Importantly, Sfp1 depletion caused an opposite transcriptional response to that of Ifh1 depletion, with Cat III RPGs being the most downregulated, and Cat I and II genes the least affected (**Figure 6C**). Remarkably, these results are fully consistent with ChEC-seq results suggesting that Sfp1 binds more strongly to Cat III than Cat I or Cat II gene promoters, with the opposite being true for Ifh1 (**Figure 5E**). It is also interesting to note that the three RPGs the least affected by Ifh1 depletion (*RPL3, RPL4A, RPL4B*) were also the most downregulated ones following Sfp1 depletion (**Figure 6D**), highlighting the complementary action of these two stress-sensitive TFs. Moreover, changes of RNAPII occupancy upon Sfp1 depletion are highly similar to those observed following TORC1 inhibition in the CARA strain, where promoter release of Ifh1 is specifically blocked (**Figure 6E**). This latter result strongly supports the idea that Sfp1 is the other factor required for coordinated repression of RPGs together with Ifh1. In this model, Ifh1 is the main regulator of Cat I and II genes but can also influence Cat III genes, whereas Sfp1 modestly affects Cat I and II genes but is the key regulator at Cat III genes. The coordinated action of these two stress-sensitive TFs is required to coordinate RPGs promoter activity upon stress. Consistent with this claim, only double depletion of Ifh1 and Sfp1 leads to a level of RPG downregulation that correlates well with what is observed following TORC1 inactivation (**Figure 6F**).

**Figure 6.**
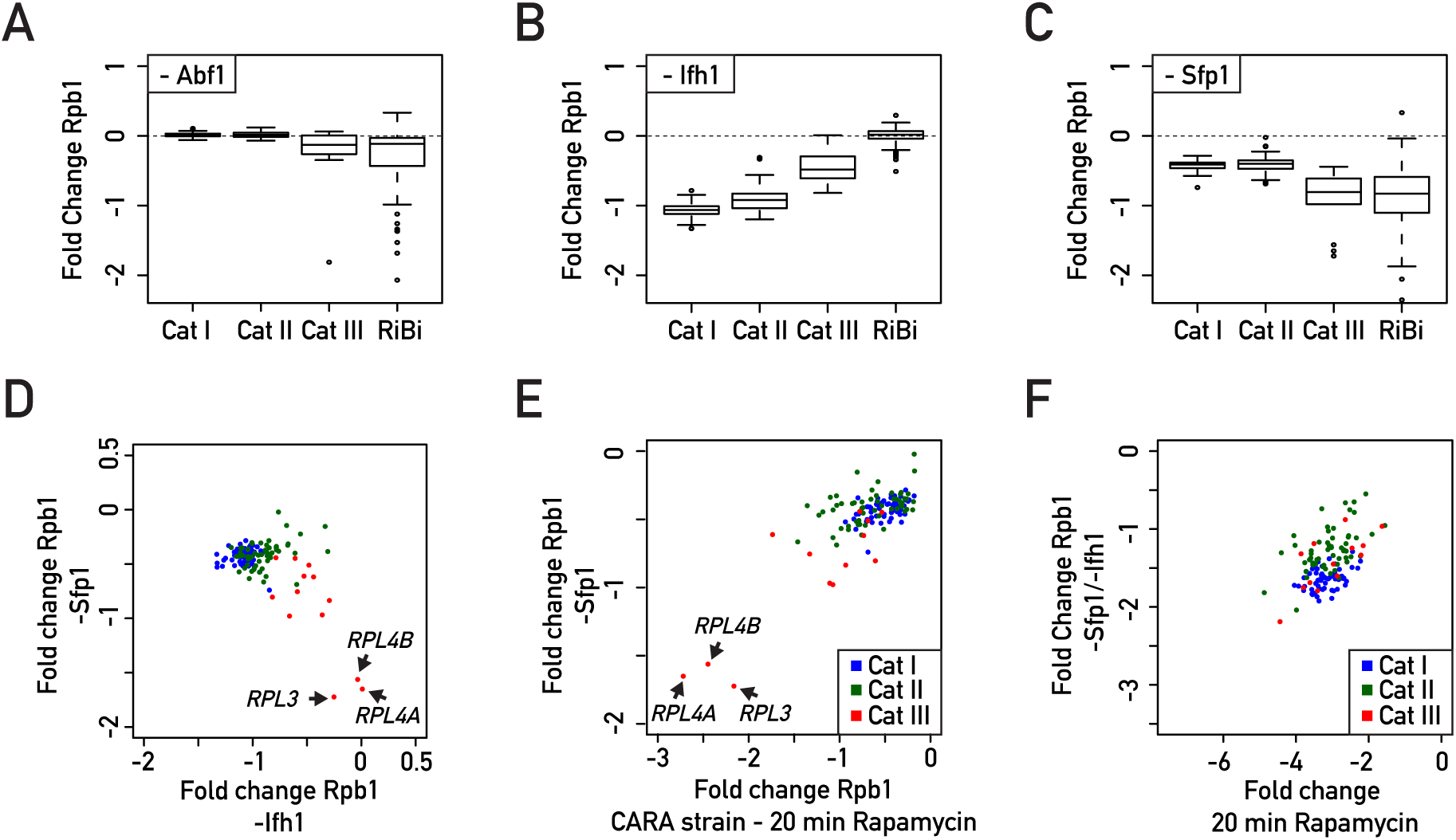
Coordinated regulation of RPGs expression is accomplished by the complementary actions of Sfp1 and Ifh1. **(A, B, C)** Box plots showing RNAPII binding change measured by Rpb1 ChIP-seq in Abf1 **(A)**, Ifh1 **(B)** or Sfp1 **(C)** nuclear-depleted cells (calculated as log2 ratio of nuclear-depleted vs. non-depleted cells) for RPGs and Ribi genes. **(D, E, F)** Scatter plots comparing RNAPII (Rpb1) binding fold change for Sfp1 nuclear-depleted cells (-Sfp1; Y-axis) versus Ifh1 nuclear-depleted cells (-Ifh1; X-axis) **(D)**; in Sfp1 nuclear-depleted cells (-Sfp1; Y-axis) versus CARA strain cells treated with Rapamycin for 20 min (CARA; x-axis) **(E)**; in double depletion of Ifh1 and Sfp1 (-Sfp1-Ifh1; Y-axis) versus WT cells treated with Rapamycin for 20 min (X-axis) **(F)**. Each dot represents a gene color-coded according to functional group as above (blue: Cat I, green: Cat II, red: Cat III).

## Discussion

Although DNA sequences features and TF binding at RGP promoters have been extensively studied using genome-wide methods in yeast (Kasahara et al., 2007; Knight et al., 2014; Reja et al., 2015; Zeevi et al., 2011), significant gaps persist in our understanding of how RPGs with heterogeneous promoter organization are coordinately regulated in response to growth and stress signals. In fact, RPG promoters that are not bound by the GRF Rap1 and which lack a duplicate copy under the control of Rap1-dependent mechanisms, such as *RPL3* and *RPL4A/B*, were thought not to be coordinately expressed with other RPGs (Reja et al., 2015). Our results demonstrate that all RPGs are regulated by the complementary action of the stress-sensitive TFs Ifh1 and Sfp1, which clarifies this apparent paradox. The results described here provide a more complete picture of the involvement of specific TFs in regulating RPG expression during stress. Our principal findings regarding the architecture of RPG promoters and their regulatory factors are summarized in a schematic form in **Figure 7**.

**Figure 7.**
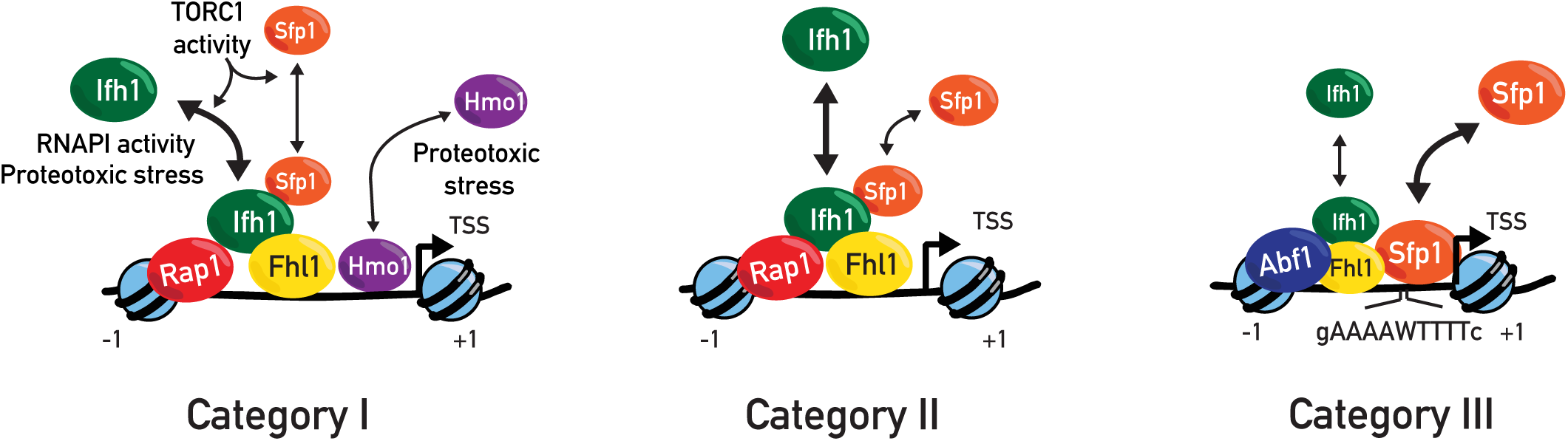
Schematic representation of proposed model of heterogeneous organization of RPGs promoters and their regulatory factors. All RPGs are regulated by two distinct, but complementary mechanisms driven by Sfp1 and Ifh1 that are required to coordinate RPG transcription upon stress. Category I and Category II RPG promoters contain the general regulatory factor Rap1 and the transcription factors Ifh1, Fhl1, Sfp1 at their promoters. Category I genes are in addition bound by the HMGB protein Hmo1. Category III promoters are bound by Abf1 (except for *RPL1A* and *RPL18B*) and are also regulated by Sfp1 and Ifh1. Ifh1 and Sfp1 release under various stress conditions downregulates RPG transcription. Ifh1 is specifically sensitive to proteotoxic stress, RNAPI activity, and TORC1 inactivation. Category III promoters are bound by the general regulatory factor Abf1 and at these RPGs Sfp1 is the key regulator. Ifh1 is the main regulator of Cat I and II but can also influence Cat III, whereas Sfp1 modestly affects Cat I and II but strongly regulates Cat III genes. The coordinated action of these two stress sensitive transcription factors are required to coordinate RPG promoter activity upon stress.

In this study, we revealed unexpected features of Hmo1 by showing its dynamic binding sensitivity upon proteotoxic stress. Interestingly, it was reported that Hmo1 transiently partitions into an aggregated protein fraction after exposure at 42°C, suggesting that the decrease in Hmo1 binding at promoters could be linked to its sequestration in an insoluble or phase-separated state (Hall et al., 2006; Reja et al., 2015; Wallace et al., 2015). Further investigation will be required to determine whether the ability of Hmo1 to transiently aggregate could be linked to some regulatory function at RPG promoters or at rDNA genes, where Hmo1 is also enriched. Nevertheless, it is becoming increasingly apparent that ribosome biogenesis is directly linked to regulation of protein homeostasis (Albert et al., 2019a; Sung et al., 2016a; Sung et al., 2016b; Tye et al., 2019). We recently demonstrated that ribosome assembly impairment in yeast triggers a rapid stress response in which RPGs are strongly downregulated and Hsf1 target genes are upregulated (Albert et al., 2019a). Moreover, we showed that this so-called Ribosome Assembly Stress Response (RASTR) is driven by orphan RP aggregation, triggering sequestration of Ifh1 in an insoluble fraction, whereas Sfp1’s activity remains unaffected during RASTR. The fact that Sfp1 activity is not affected by RASTR raises the issue of over-accumulation and aggregation of RPs encoded by genes controlled by Sfp1, such as *RPL4A/B* and *RPL3*, which are weakly affected by Ifh1 depletion or during RASTR. Interestingly, both Rpl4 and Rpl3 belong to a restrictive group of RPs with a dedicated chaperone, suggesting that the latter can compensate for the absence of Ifh1-dependent downregulation upon stress (Pillet et al., 2017). On the other hand, protein quality control (PQC) mechanisms may contribute to prevent aggregation of other unassembled RPs. Accordingly, in strains where the excess ribosomal protein quality control (ERISQ) pathway is ablated (e.g. *tom1Δ* cells), we showed that Crf1 expression becomes toxic, indicating that regulation of RPG transcription is important to maintain protein homeostasis.

Over twenty years ago Laferté et al. (2006) described a yeast strain expressing a version of RNAPI that remains constitutively active under stress, which led to the notion of the central role of this enzyme in coordinating stoichiometric levels of ribosome components (Chedin et al., 2007). Similarly, we constructed here a strain (overexpressing *CRF1*) that prevents the proper regulation of RPG expression, which normally accounts for about 50% of RNAPII initiation events in growing cells (Warner, 1999). This strain represents a promising tool to understand the importance of modulation of RPG expression during the cell cycle, stress or meiosis. For example, several studies have shown that splicing machinery is present in limiting amounts, suggesting that the splicing process may also be modulated by changing the amount of pre-mRNA substrate containing introns (Munding et al., 2013). Given that RPGs produce the major part of spliced pre-mRNA and that endoplasmic reticulum stress or meiosis both involve downregulation of RPGs and splicing of stress- or meiosis-specific genes, respectively (Kawahara et al., 1997; Munding et al., 2010), it could be interesting to challenge the importance of RPG downregulation in these processes by using this *CRF1*-overexpressing strain. It could be also interesting to determine to what extent the maintenance of a high pool of RPG mRNAs upon stress could prevent proper translation of mRNAs derived from stress genes that need to be rapidly transcribed and translated in these conditions.

It might seem at first that a simpler system in which all RPGs would be controlled by a common mechanism would make more evolutionary sense. However, after the whole genome duplication preceding the emergence of *Saccharomyces cerevisiae*, most RPG paralogues were actually conserved, despite the fact that nearly 90% of all duplicated genes were eliminated. This could suggest that a higher level of complexity in RPG regulation provides a selective advantage (Coulombe-Huntington and Xia, 2017; Wapinski et al., 2007). It is tempting to hypothesize that the differential RPG regulation described in this study will also increase the ability of cells to respond to varying growth conditions. Indeed, differential RPG promoter regulation could provide plasticity to RP paralogue production, which might in turn favor functional differences in specialized ribosomes (Dinman, 2016; Kondrashov et al., 2011; Mauro and Edelman, 2002). It was also proposed that Abf1 binding at RPG promoters may help in rapid resumption of transcription after stress (Fermi et al., 2016). This differential RPG regulation could contribute to regulate specifically the expression of pleiotropic RPGs, which have other roles, in addition to their original ribosomal function (Warner and McIntosh, 2009). In this sense, it is interesting to note that genes from Cat III, which are highly sensitive to Sfp1, are enriched in those encoding extra-ribosomal functions (Reja et al., 2015). The combination of intrinsic DNA features encoded in the promoter and the ability to modulate the recruitment of a different set of TFs described in this study are probably the major determinants of RPG promoter activity in budding yeast. Pre-mRNA splicing, mRNA decay and protein turnover provide additional layers of complexity to achieve a balanced production of ribosome components that allows for efficient ribosome assembly while avoiding the proteotoxicity of unassembled RPs.

In metazoan cells, transcription of rDNA genes is relatively well studied, and its general properties are highly conserved (Grummt, 2013). However, the general features of RPG expression are much less well understood compared to yeast. Given that the defect of expression of some RPs leads to a heterogeneous class of diseases known as ribosomopathies, and that it was reported that RPGs exhibit tissue or disease specific expression patterns in mammalian cells (Guimaraes and Zavolan, 2016), future studies should seek to identify mechanisms orchestrating RPG expression in metazoan cells and organisms.

## Material and methods

### Yeast strains and growth conditions

The experiments presented in this study were performed using the budding yeast *Saccharomyces cerevisiae*. A complete list of strains and plasmids used is provided in Tables S7 and S8, respectively. Strains were generated by genomic integration of tagging or disruption cassettes as described (Longtine et al., 1998; Rigaut et al., 1999). Yeast cells were grown in YPAD medium (1% yeast extract, 2% peptone, 20 mg/mL adenine sulfate, 2% glucose) at 30°C overnight and then the cultures were diluted to OD_600_ = 0.1. Most experiments were performed with exponential phase cells harvested between OD_600_ 0.4 and 0.6, unless otherwise indicated.

For conditional depletion of target proteins briefly, cells were grown to log-phase (OD_600_ = 0.3–0.4) and anchor-away of FRB-tagged proteins was induced by treating cells with rapamycin (1 mg/ml of stock solution resuspended in 90% ethanol, 10% Tween-20) to 1 µg/mL final concentration for 20 min and 60 min before collection (Haruki et al., 2008). Rapid depletion of AID-tagged proteins was induced by the addition of 3-indoloacetic acid (auxin) at 500 mM final concentration. For heat stress experiments, cells were incubated in complete medium until log phase at 30°C, then the samples were briefly centrifuged, and pellets were resuspended in pre-warmed medium at the indicated temperatures and incubated for 5 min. The cells were collected following various treatments at the indicated time points before crosslinking. Arrest of translation was induced by adding cycloheximide to a final concentration of 25 µg/ml. Cells were treated with diazaborine to a final concentration of 50 µg/ml and with tunicamycin to a final concentration of 1 µg/ml.

### Growth assays

Tenfold serial dilutions of log-phase growing cells (OD_600_ = 0.3) were spotted on plates containing complete medium (YPAD) or synthetic selective medium at indicated temperatures. Plates were imaged following 24h and 48h incubations.

### ChIP and ChIP-Seq

Yeast cultures of 50 mL in complete medium were collected at OD_600_ = 0.4–0.6 for each condition. The cells were crosslinked with 1% formaldehyde for 10 min and quenched by adding 125 mM glycine for 5 min at room temperature. Cells were then washed with ice-cold HBS and resuspended in 0.6 mL of ChIP lysis buffer (50 mM HEPES-Na pH:7.5, 140 mM NaCl, 1 mM EDTA, 1% NP-40, 0.1% sodium deoxycholate) supplemented with 1 mM PMSF and 1x protease inhibitor cocktail (Roche). The cells were broken using Zirconia/Silica beads (BioSpec) and lysates were centrifuged at 13,000 rpm for 30 min at 4°C. Following centrifugation, the pellets were resuspended in 300 μL ChIP lysis buffer containing 1 mM PMSF and sonicated for 15 min (30s ON - 60s OFF) in a Bioruptor (Diagenode). The lysates were centrifuged at 7000 rpm for 15 min at 4°C, following which primary antibodies were added to the supernatant and incubated for 1 hr at 4°C. ChIP was performed using the following antibodies: for RNAPII (Abcam ab5131), for Myc, (homemade mouse antibody, clone 9E10); for Hmo1 (produced by Pocono Farm; rabbit). Magnetic beads were washed three times with PBS containing 0.5% BSA and added to the lysates (30 μL of beads/300 mL of cell lysate). The samples were incubated for 2 hr at 4°C. The beads were washed twice with AT1 buffer (50 mM HEPES-Na pH: 7.5, 140 mM NaCl, 1 mM EDTA, 0.03% SDS), once with AT2 buffer (50 mM HEPES-Na pH: 7.5, 1 M NaCl, 1 mM EDTA), once with AT3 buffer (20 mM Tris-Cl pH: 7.5, 250 mM LiCl, 1 mM EDTA, 0.5% NP-40, 0.5% sodium deoxycholate) and twice with TE buffer. The chromatin was eluted from the beads by resuspension in TE containing 1% SDS and incubation at 65°C for 10 min. The eluate was transferred to new Eppendorf tubes and incubated overnight at 65°C to reverse the crosslinks. The DNAs were purified using the MinElute PCR Purification Kit (Qiagen). DNA libraries were prepared using TruSeq ChIP Sample Preparation Kit (Illumina) according to manufacturer’s specifications. The libraries were sequenced using an Illumina HiSeq 2500 platform at the Institute of Genetics and Genomics of Geneva (iGE3; http://www.ige3.unige.ch/genomics-platform.php) and the reads were mapped to the sacCer3 genome assembly using HTSStation60 (read densities were calculated using shift: 100bp, extension: 50bp). All densities were normalized to 10M reads.

For RNAPII, the signal was quantified for each gene between the transcription start site (TSS) and transcription termination site (TTS). To quantify ChIP-seq signals for each promoter, a ratio between the total number of reads from each sample in a 400 bp region upstream the TSS (Jiang and Pugh, 2009) for each ORF and the total number of reads from the same region were obtained with mock IP of the control untagged strain. To compare depleted versus non-depleted cells, we divided the signal from the +auxin and/or +rapamycin samples by the signal from the -auxin and/or -rapamycin (vehicle) samples and log2 transformed this value. All data from publicly available databases were mapped using HTS Station (http://htsstation.epfl.ch; (David et al., 2014)). For ChIP-seq experiments, cross-linked chromatin obtained from fission yeast *Schizosaccharomyces pombe* was used as a spike-in control (Bruzzone et al., 2018). Values of ChIP-seq signal for each gene are reported in Supplementary Tables 2,3,4,6.

### ChEC-seq

ChEC-seq experiments were performed as described (Zentner et al., 2015). ChEC-seq data from Albert et al. (2019b) were used to determine sites of Sfp1 and Ifh1 binding. A strain expressing ‘free’ MNase under control of the *REB1* promoter was used as control. Briefly, cells were washed twice with buffer A (15 mM Tris 7.5, 80 mM KCl, 0.1 mM EGTA, 0.2 mM spermine, 0.5 mM spermidine, 1xRoche EDTA-free mini protease inhibitors, 1 mM PMSF) and resuspended in 200 μL of buffer A containing 0.1% digitonin. The cells were incubated at 30°C for 5 min. MNase action was then induced by addition of CaCl_2_ to 5 mM and stopped at the indicated time points with EGTA at a final concentration of 50 mM. DNA was purified using MasterPure Yeast DNA purification Kit (Epicentre) according to the manufacturer’s specifications. Large DNA fragments were removed by a 5-min incubation with 2.5x volume of AMPure beads (Agencourt) after which the supernatant was kept, and MNase-digested DNA was precipitated using isopropanol.

Libraries were prepared using NEB Next kit (New England Biolabs) according to the manufacturer’s protocol. Prior to PCR amplification of the libraries, small DNA fragments were selected with a 5-minute incubation using 0.9x volume of the AMPure beads after which the supernatant was kept and incubated with the same volume of beads as before for another 5 min. DNA was eluted with 0.1x TE after washing the beads with 80% ethanol and PCR was performed. Adaptor dimers were removed with a 5 min incubation using 0.8x volume of the AMPure beads after which the supernatant was kept and incubated with 0.3x volume of the beads. The beads were then washed twice with 80% ethanol and DNA was eluted using 0.1x TE. The quality of the libraries was verified by running an aliquot on a 2% agarose gel. Libraries were sequenced using a HiSeq 2500 machine in single-end mode. Reads were extended by the read length. Reads were mapped to the genome (sacCer3 assembly) using HTSStation, and the position of the 5′-most base of each read was used as the position of the MNase cut site. All densities were normalized to 10M reads. For peak analysis, we used the signal obtained for Sfp1-MNase or Ifh1-MNase after 30 seconds or 2 minutes 30 seconds of CaCl_2_ treatment, respectively, normalized by dividing it by the signal obtained for free MNase after 20 min of treatment. Values of ChEC-seq signal at each promoter are reported in Supplementary Table 5.

### Heat maps, plots and statistics

In all of the box plots, the box shows the 25th–75th percentile, whiskers show the 10th–90th percentile, and dots show the 5th and 95th percentiles. Statistical significance of difference between groups was evaluated using the Mann-Whitney rank sum test.

### Data and software availability

All deep sequencing data sets have been submitted to the NCBI Gene Expression Omnibus under accession code GSE XX

## Acknowledgments

We thank to all members of the Shore lab for comments and discussions throughout the course of this work; Mylène Docquier and the Genomics Platform of iGE3 at the University of Geneva (https://ige3.genomics.unige.ch/) for high-throughput sequencing services; Prof. Helmut Bergler (Karl-Franzens-Universität, Graz, Austria) for his generous gift of diazaborine; Nicolas Roggli for expert assistance with data presentation and artwork. B.A. acknowledges support from a long-term EMBO postdoctoral fellowship in the early phases of this work and D.S. acknowledges funding from the Swiss National Science Foundation (grant number 31003A_170153) and the Republic and Canton of Geneva.

## Author Contributions

B.A., S.Z and D.S. conceived the study. B.A., S.Z, M.P.R and D.D. performed most experiments and analysed the results together with D.S. B.A, S.Z, and D.S. wrote the manuscript.

---

**Supplemental Figure 1.**
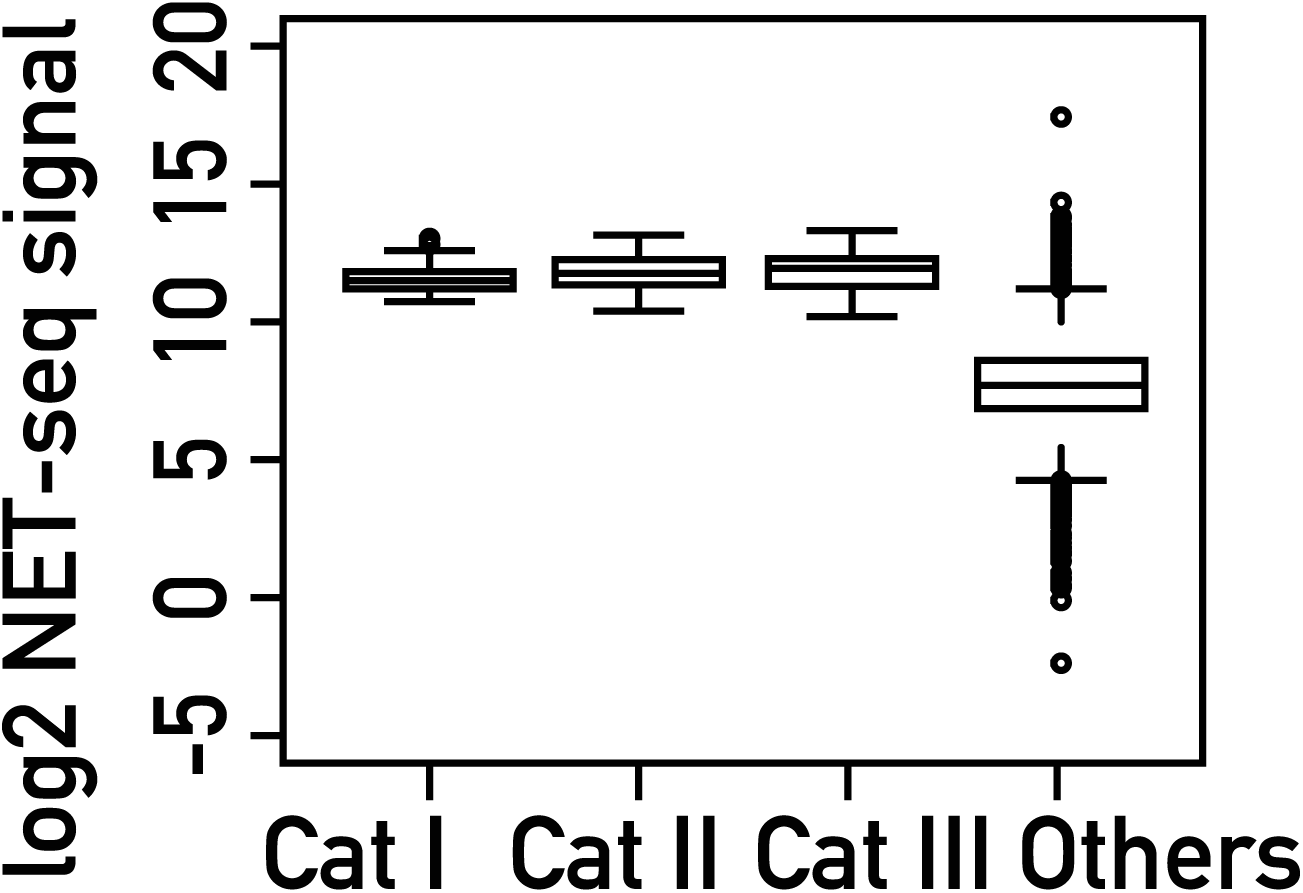
Box plots showing the transcription rate measured by nascent elongating transcript sequencing (NET-seq; (Churchman and Weissman, 2011)) for categories I, II and III of RPGs, and all other genes.

**Supplemental Figure 6.**
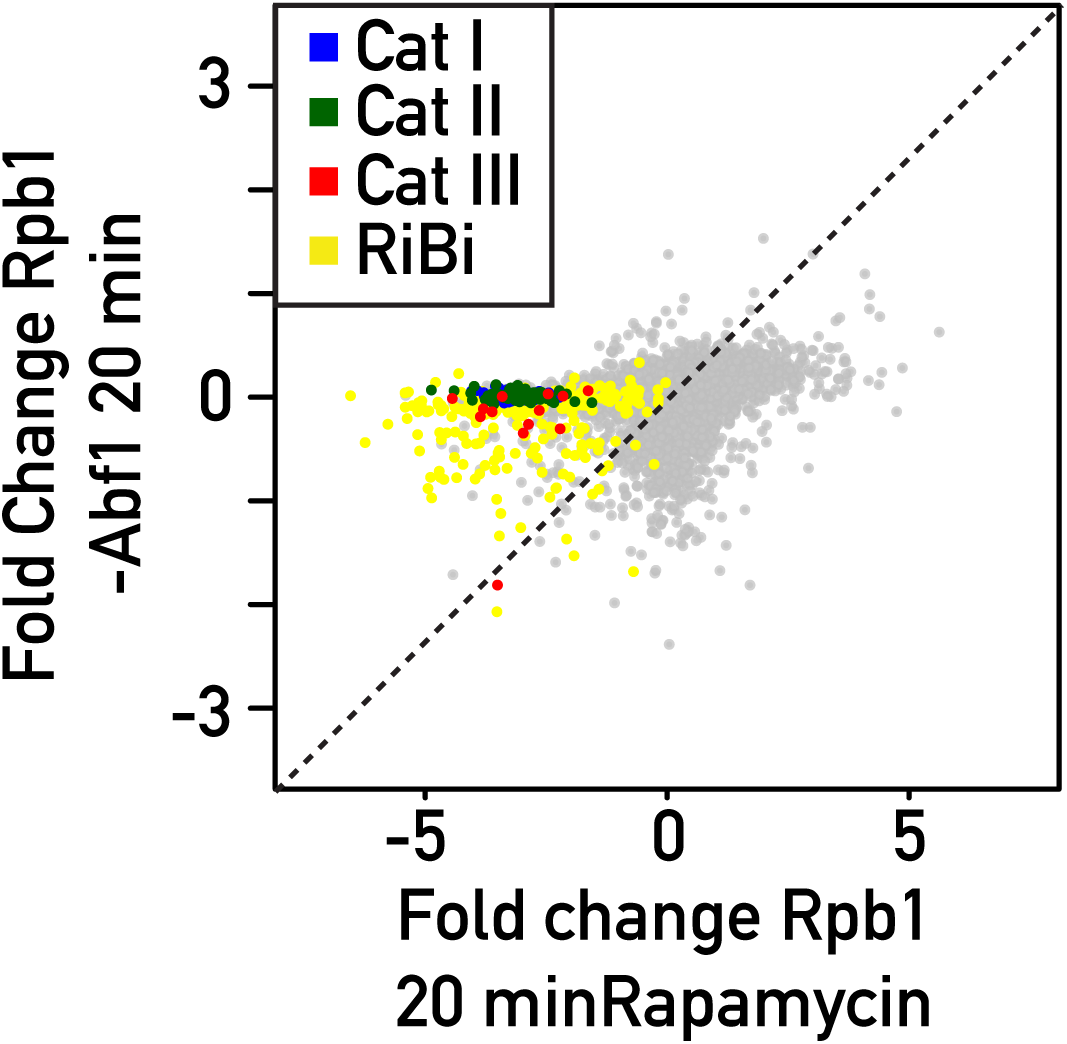
RNAPII ChIP-seq in Rpa135 nuclear-depleted cells (Rpa135-depleted: Y-axis) versus non-depleted cells (vehicle treatment: X-axis). Bottom panels display the corresponding box plots for the four indicated gene categories.

## Notes

### Competing Interest Statement

The authors have declared no competing interest.

